# A comprehensive phylogenetic analysis of the serpin superfamily

**DOI:** 10.1101/2020.09.09.289108

**Authors:** Matthew A. Spence, Matthew D. Mortimer, Ashley M. Buckle, Bui Quang Minh, Colin J. Jackson

**Affiliations:** Research School of Chemistry, Australian National University, ACT 2601, Australia; Biomedicine Discovery Institute, Department of Biochemistry and Molecular Biology, Monash University, Victoria, 3800, Australia; Research School of Computer Science and Research School of Biology, Australian National University, ACT 2601, Australia

## Abstract

Serine protease inhibitors (serpins) are found in all kingdoms of life and play essential roles in multiple physiological processes. Owing to the diversity of the superfamily, phylogenetic analysis is challenging and prokaryotic serpins have been speculated to have been acquired from Metazoa through horizontal gene transfer (HGT) due to their unexpectedly high homology. Here we have leveraged a structural alignment of diverse serpins to generate a comprehensive 6000-sequence phylogeny that encompasses serpins from all kingdoms of life. We show that in addition to a central “hub” of highly conserved serpins, there has been extensive diversification of the superfamily into many novel functional clades. Our analysis indicates that the hub proteins are ancient and are similar because of convergent evolution, rather than the alternative hypothesis of HGT. This work clarifies longstanding questions in the evolution of serpins and provides new directions for research in the field of serpin biology.

## Introduction

The <u>ser</u>ine <u>p</u>rotease <u>in</u>hibitor (serpin) superfamily is the largest group of protease inhibitors in nature (Gettins 2003). Serpins have been identified throughout all kingdoms of life: animals, plants, fungi, protists, archaea and bacteria (Law et al. 2006). It is notable that eukaryotic serpins are significantly more abundant than their prokaryotic counterparts, which are found in only a few lineages of bacteria and archaea (Irving et al. 2002). Indeed, many eukaryotic serpins have extremely well understood physiological roles (for example, alpha-1 antitrypsin, A1AT), whereas the functions of prokaryotic serpins are relatively enigmatic. This has contributed to controversy regarding the progenitor of prokaryotic serpins and the evolutionary origins of the superfamily (Irving et al. 2002; Roberts et al. 2004; Kantyka et al. 2010; Ivanov et al. 2006; Goulas et al. 2017). The ubiquitous presence of serpins in plant and metazoan biology has led to the hypothesis that serpins are a relatively young superfamily that emerged in the last common ancestor of eukaryotes (Irving et al. 2000). Under this model of serpin evolution, prokaryotes acquired serpins *via* inter-kingdom horizontal gene transfer from a eukaryote (Irving et al. 2002).

Canonical, inhibitory members of the superfamily have evolved to exploit the energetic difference between two physiologically relevant conformations: in the metastable stressed (native) state, a solvent exposed and flexible reactive centre loop (RCL) protrudes from the central β-sheet of the protein (the A-sheet), resembling the typical unstructured peptide substrates of target proteases. Cleavage of the RCL at the scissile bond by an attacking protease induces a conformational change to the relaxed, lower energy cleaved state, in which the cleaved RCL is inserted as an additional strand of the A-sheet. The stressed (S) to relaxed (R) conformational transition kinetically traps the covalent serpin-protease complex, irreversibly inhibiting the attacking protease by distorting the active site residues. Serpins thus function as irreversible, suicide inhibitors. The specificity of serpins towards their cognate protease targets is determined by the amino-acid sequence of the RCL (Huntington 2011), the complex interplay between dynamics in and around the RCL, and local electrostatics (Marijanovic et al. 2019), as well as auxiliary exosites that stabilise the initial non-covalent serpin-protease association complex prior to RCL cleavage (Gettins and Olson 2009). The functional dependence on the relative energies of the R and S states makes the serpin inhibitory mechanism particularly-well suited to allosteric regulation by cofactors; stabilization or destabilization of one conformation by a ligand can alter this energetic balance and modulate serpin function. The archetype of allosteric modulation in the serpins is the regulation of antithrombin by the binding of heparin to an exosite, which accelerates inhibition by acting as a template for both serpin and protease (Johnson et al. 2006, Figure 1).

**Figure 1.**
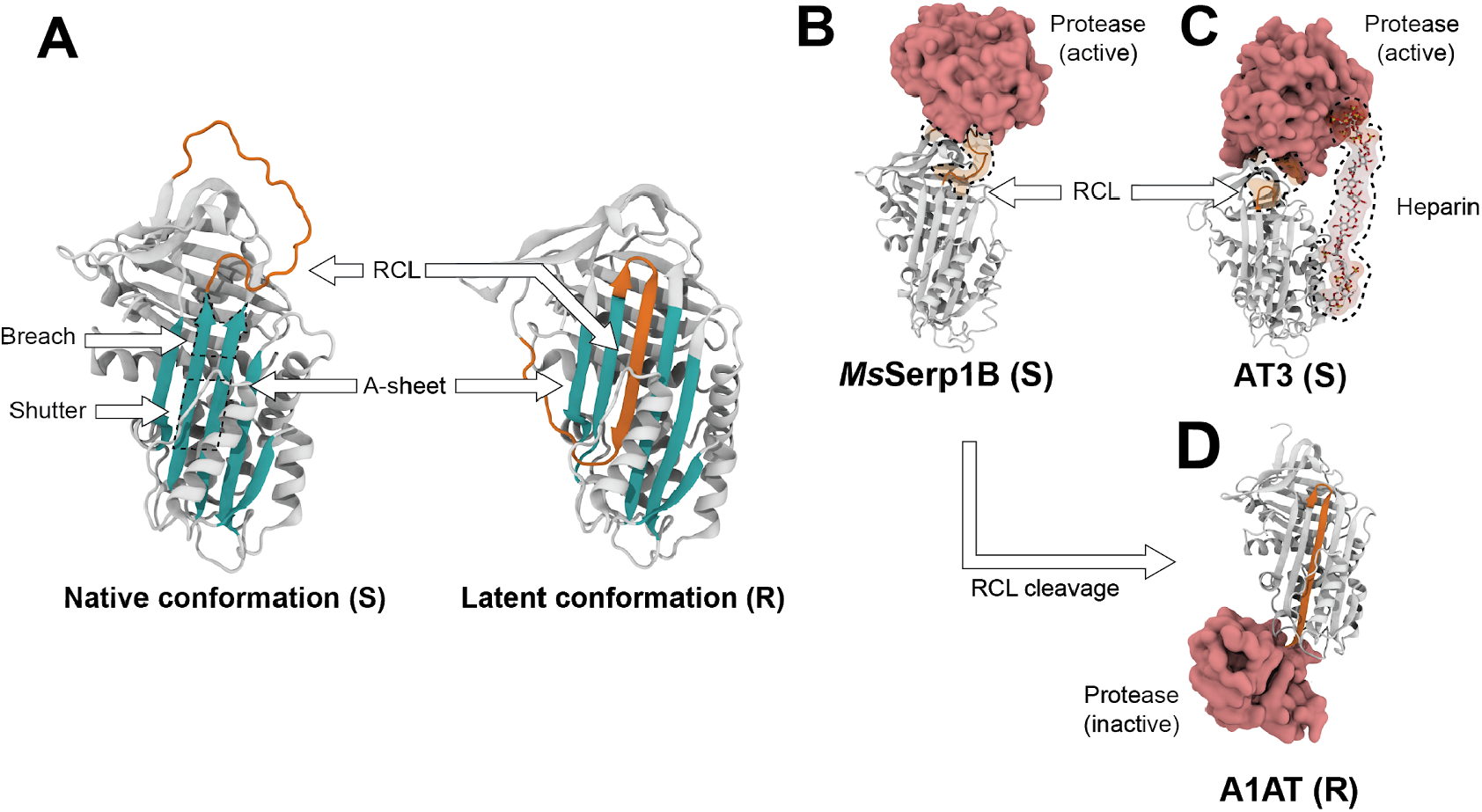
Serpin fold and mechanistic diversity. (S) and (R) denote stressed and relaxed conformations, respectively **A**. Cartoon representation of the S to R state conformational transition, in alpha-1-antitrypsin (A1AT). The RCL in both native (exposed) and latent (inserted) conformations is highlighted, among other key structural components of the serpin fold, such as the breach, shutter and A-sheet (PDBs: native stressed conformation 3NE4, latent relaxed conformation 1IZ2). **B**. non-covalent Michaelis complex between *Manduca sexta* serpin1B (*Ms*Serp1B) and active trypsin protease (PDB: 1K9O). **C**. Ternary michaelis complex between human antithrombin III (AT3) with heparin templated active thrombin (PDB: 1TB6). **D**. covalent complex between cleaved A1AT and inhibited trypsin, post RCL cleavage (PDB: 1EZX).

While the canonical serine protease inhibitor function is the dominant function within the superfamily and has been characterized in great detail, it is clear that the functional diversity of the serpins extends well beyond serine protease inhibition. For example, cross-class and papain-like cysteine protease inhibitors have been documented (Schick et al. 1998; Kantyka et al. 2011; Guo et al. 2015; Zhou et al. 1997). Many members of the serpin superfamily have also been shown to possess non-inhibitory functions. In these instances, the non-inhibitory functions typically exploit the conformational transition from S to R states, such as the corticosteroid and thyroxine binding globulins that transport steroid hormones in higher eukaryotes *via* differential affinity for steroid ligands in the R and S forms (Zhou et al. 2008; Zhou et al. 2006).

Understanding the molecular basis of the functional divergence within protein superfamilies can provide insight into the biophysical properties that permit or constrain functional radiation in protein evolution and reveal the determinants of novel phenotypes, as well as the selective pressures that compel organisms to acquire them. Modeling the evolution of protein superfamilies is often challenging due to poor phylogenetic signal, extensive sequence diversity and large sequence datasets that have been diverging over geological timescales. This is particularly true of serpins; the superfamily’s broad distribution and extensive sequence divergence have limited previous phylogenetic analyses to specific serpin families and taxonomic groups, or obscured the topology of deep branches in the serpin phylogeny (Irving et al. 2000; Heit et al. 2013). Additionally, the number and diversity of serpin sequences belonging to prokaryotes has continued to expand since the advent of the genomic age, further clouding the evolutionary history of serpins. Here, we leverage structural information from the consensus serpin fold to perform comprehensive phylogenetic analysis of the serpin superfamily. This phylogeny provides new insight into the evolution of serpins across the various kingdoms of life and has generated new hypotheses relating to the possible biological function of a number of previously sparsely characterized clades and the origins of the superfamily.

## Results and Discussion

### A global perspective on serpin sequence space from sequence similarity network (SSN) analysis

To investigate global serpin sequence diversity, we generated an unbiased and non-redundant dataset of 18233 serpin proteins and putative gene products. This dataset was then analysed as a sequence similarity network (SSN), in which nodes represent sequences (or redundant clusters of sequences) and edges connecting nodes embody similarity scores above an arbitrary similarity threshold (Figure 2). Unlike phylogenetic analysis, an SSN can only convey information relating to the similarity between sequences in a dataset and does not explicitly model sequence evolution or phylogenetic relatedness (Atkinson et al. 2009; Copp et al. 2018); however, SSNs have been profoundly insightful and have proven to be useful in revealing sequence-function relationships in a variety of protein superfamilies without the technical and computational limitations and difficulties of full phylogenetic inference (Akiva et al. 2017; Ahmed et al. 2015; Wichelecki et al. 2015; Baier and Tokuriki 2014; Akiva et al. 2013).

**Figure 2.**
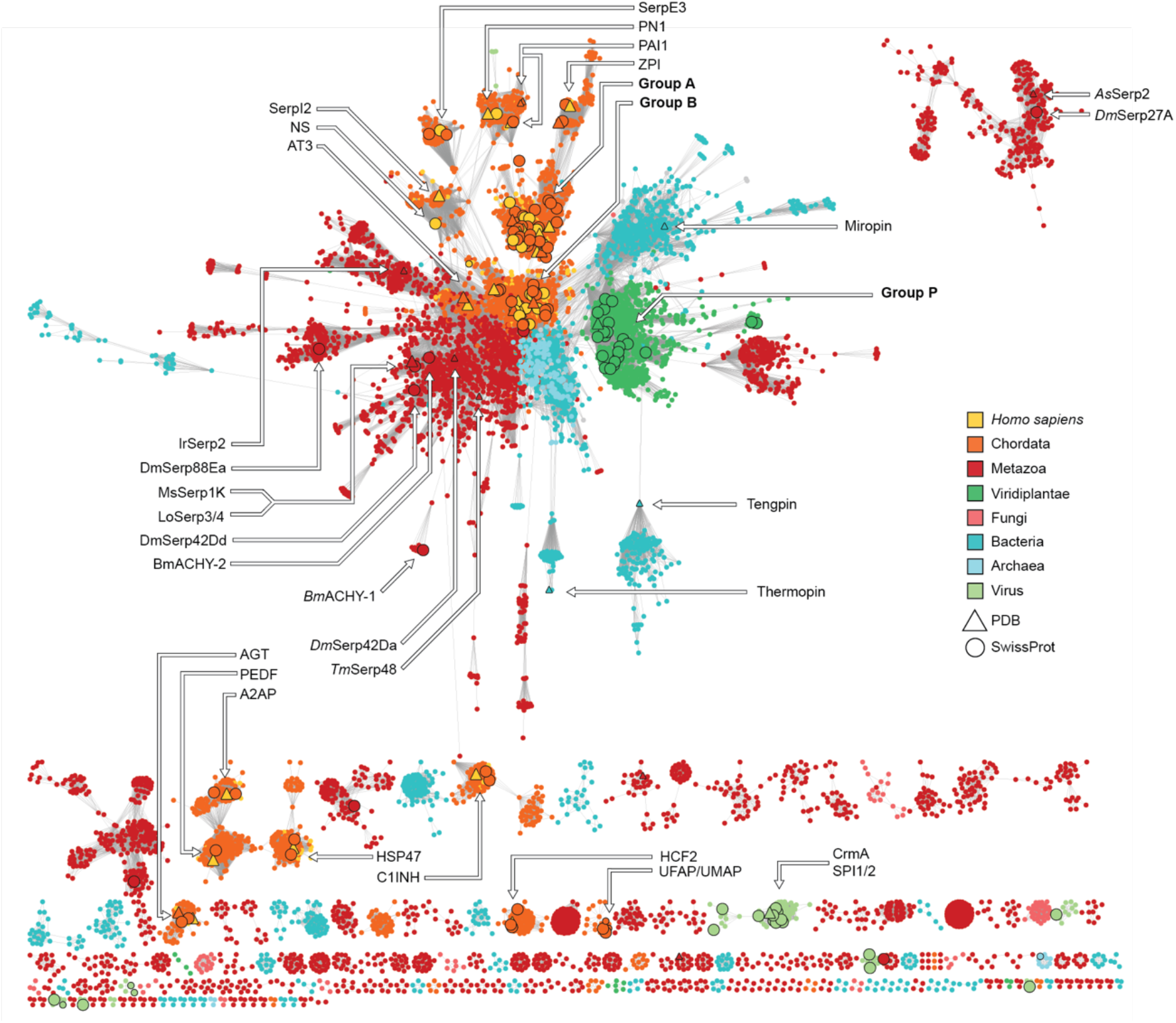
A non-redundant sequence similarity network of the serpin superfamily. Nodes represent serpin sequences, or clusters of serpin sequences that share 75% pairwise sequence identity. Edges connect sequences that share >40% pairwise sequence identity. Nodes are coloured by the taxonomic distribution of organisms that they belong to: nodes that represent chordate and *H. sapiens* serpins are distinguished at the level of phylum and species for clarity. Sequences with reviewed annotations and solved structures are represented by enlarged circular and triangular nodes with black borders, respectively.

In the global view of serpin sequence space generated by SSN analysis, 10123 nodes represent 18233 amino acid sequences (sequences with >75% similarity are collapsed into a single node) that constitute the serpin protein superfamily (pfam: PF0079) and edges connect nodes that share greater than 40% pairwise sequence identity. We employed the ‘divide and conquer’ workflow of Akiva and Copp (Akiva et al. 2017; Copp et al. 2018) in conjunction with simulated Markov network clustering (MCL) (Morris et al. 2011) to define 481 discrete and highly interconnected clusters of sequences that are expected to be functionally distinct from one another. Indeed, this sequence clustering workflow distinguished annotated subgroups of highly homogenous Group A and Group B serpins from one another exclusively on the basis of SSN topology (Supplementary Figure 1), indicating that our classification criteria could effectively differentiate functionally distinct serpin families.

The serpin superfamily SSN shares many characteristics with those of other protein superfamilies and biological systems. The network is approximately scale-free (i.e. the edge distribution follows a power-law), as is often typical of biological and evolutionary networks, and serpins from closely related taxa tend to cluster together. Most sequences belong to a central connected component that exhibits a hub-and-spoke topology, in which divergent groups radially descend from a central cluster of sequences that we hereafter refer to as the ‘hub’. The central component includes chordate Groups A, B, C, E and I, arthropod Group K and plant Group P, according to the classification scheme defined by (Irving et al. 2000), as well as the majority of unclassified prokaryotic serpins (Figure 2). Many serpins that have diverged from canonical inhibitory functions, such as uterine associated serpins (UMAP, UFAP) and HSP47 are disconnected from the central network component, reflecting their functional divergence (Ing and Roberts 1989; Widmer et al. 2012).

### The serpin network hub consists of similar chordate and prokaryotic serpins

The most prominent topological feature of the serpin SSN is the presence of a heterologous hub group that other clustered sequences radiate from. Hub groups are observed in other protein SSNs and sequences that occupy hub-like positions in SSNs generally share broad similarity across sequence space, because of this, hub-like sequences tend to represent the most consensus-or ancestral-like contemporary sequences (Akiva et al. 2017). The peculiarity of the serpin SSN hub is that it is composed of serpins that are highly similar to each other, yet belong to taxa that are distantly related. This is particularly evidenced by the high similarity shared between chordate Group B serpins with archaeal and bacterial serpins. On the premise of sequence similarity alone, chordate Group B serpins appear to be more closely related to almost all groups of prokaryotic serpins than to other members of the chordate serpin lineage that they are known *a priori* to belong to (Heit et al. 2013; Irving et al. 2000), such as chordate Groups H, D and even other members within Group B (UMAP,UFAP).

The incongruence of microbial and chordate serpins sharing significant homology has led to the hypothesis that prokaryotic serpins were acquired *via* a rare inter-kingdom horizontal gene transfer event recently in serpin evolution (herein referred to as the HGT hypothesis) (Irving et al. 2002; Roberts et al. 2004; Kantyka et al. 2010; Ivanov et al. 2006; Goulas et al. 2017). One underlying assumption of the HGT hypothesis is the existence of a viable mechanism for genetic transfer between a chordate and recipient prokaryote and a clear selective advantage provided by the transferred genetic material (Koonin et al. 2001;Ochman et al. 2000). This assumption may be valid when considering exclusively commensal and pathogenic prokaryotes that co-exist with chordate hosts, such as the prokaryotic serpin miropin from the opportunistic pathogen *Tannerella Forsythia;* however, the HGT hypothesis is not supported by all of the available data and is inconsistent with the full breadth of free-living prokaryotes (such as halophilic archaea belonging to *Haloferax, Natrialba*, thermophilic archaea belonging to *Thermococcus*, psychrophilic bacteria belonging to *Psychorbacter*, free-living soil bacteria belonging to *Sporangium, Chondromyces* and free-living marine sediment bacteria belonging to *Beggiatoa;* Supplementary Figure 2) that encode chordate-like serpins. Indeed, most of the prokaryotic hub serpins belong to marine and soil bacteria, sulfur and methane metabolizing prokaryotes and environmental extremophiles (with the exception of few belonging to *Elusimicrobia spp*.) and are hence unlikely to play a role in pathogenicity or host-microbe interaction, as in other clusters of prokaryotic serpins (such as those that include miropin and serpins from known commensals and pathogens). Almost all of the hub-like prokaryotic serpins do share the RCL-hinge sequence motif with known inhibitory serpins (P17 - P9: ExGTEAAAA, x:E/K/R) (Hopkins et al. 1993; Irving et al. 2000), indicating that they are competent protease inhibitors that can undergo the stressed-to-relaxed conformational transition.

We hypothesize that the hub-like prokaryotic serpins occupy primitive, intracellular housekeeping roles, such as controlling cytoplasmic proteolysis and likely resemble ancestral inhibitory serpins. Indeed, the two archaeal serpins within the hub that have experimental annotations (Pnserpin from *Pyrobaculum neutrophilum* and Tkserpin from *Thermococcus kodakaraiensis*) are both potent protease inhibitors that can effectively inhibit endogenous subtilisin and chymotrypsin-like proteases at the extreme temperatures at which the hyperthermophilic organisms reside (Zhang et al. 2017; Tanaka et al. 2011).

### Much of serpin sequence space remains unexplored

There are many other well-defined sequence clusters beyond the central hub and connected component in the serpin SSN, such as those belonging to angiotensinogen, heat-shock protein 47 (HSP47),UFAP/UMAP, C1 inhibitor (C1INH) and heparin cofactor 2 (HCF2), which have been functionally characterized through comprehensive work on chordate serpins. However, the majority of serpin sequence space lacks annotation. Of the 481 clusters we identified (minimal criteria of at least 4 unique sequences in a cluster), only 13.5% have representative members with robust, experimental annotation or structural representation. Most of these belong to chordate serpins and despite accounting for the majority of the serpin superfamily’s total diversity, only 3.5% of non-chordate metazoan and 5.2% of prokaryotic serpin clusters larger than 4 unique sequences have at least a single member with experimental annotation. Among the groups that lack any annotation are 40 sequence clusters with comparable size (>20 sequences) to the major Chordate serpin subfamilies that are likely to be functionally distinct. Additionally, the current serpin classification scheme devised from the first published phylogeny of the serpin superfamily includes only a fraction of the sequence diversity that we find here (Irving et al. 2000). There is also no robust, cladistic classification scheme for bacterial and archaeal serpins, emphasizing a broad necessity within the field to experimentally investigate diverse, non-chordate serpins and justifying the need for updated phylogenetic studies on the serpin superfamily.

### A unified phylogeny of the serpin superfamily

To study the evolution of serpins, we performed a comprehensive, superfamily-wide phylogenetic analysis. Whereas SSNs exclusively provide insight on the global similarities shared between protein sequences, full phylogenetic analysis can deconvolute the evolutionary topologies that relate extant sequences to one another at the expense of computational burden. Large-scale phylogenetic inference of full protein superfamilies is technically challenging; homology is often difficult to detect between distantly related members of a superfamily and the difficulty of attaining an accurate sequence alignment is often a barrier to phylogenetic inference.

To overcome this, we devised a workflow to align serpin sequences within our dataset based on structural information derived from the conserved serpin fold. Because a protein structure contains more evolutionary information than an amino-acid sequence alone (Shakhnovich et al. 2005), we were able to align a diverse subset of 750 representative serpin sequences (selected as representatives of 750 clusters from the SSN at a 52.5% sequence similarity threshold) to a hidden Markov model (HMM) trained on the distributions of residues observed at each position in 39 structurally aligned and evolutionarily diverse serpin crystal-structures (Supplementary Figure 3). Using this structure-guided sequence alignment as a seed for a full sequence dataset produced an alignment of 6,000 unique serpin amino-acid sequences that were robustly aligned at all positions of the consensus serpin fold that are crystallographically resolved. This meant that the RCL (C-terminal of the conserved hinge motif), N-and C-terminal extensions, as well as non-conserved insertions throughout the serpin scaffold that are generally not resolved in or are absent from solved serpin structures were omitted from the alignment. Hypervariability among these sites indicates that they may not be related by homology and should not be included in the alignment regardless, as in previous phylogenetic analyses of the serpins (Irving et al. 2000).

**Figure 3.**
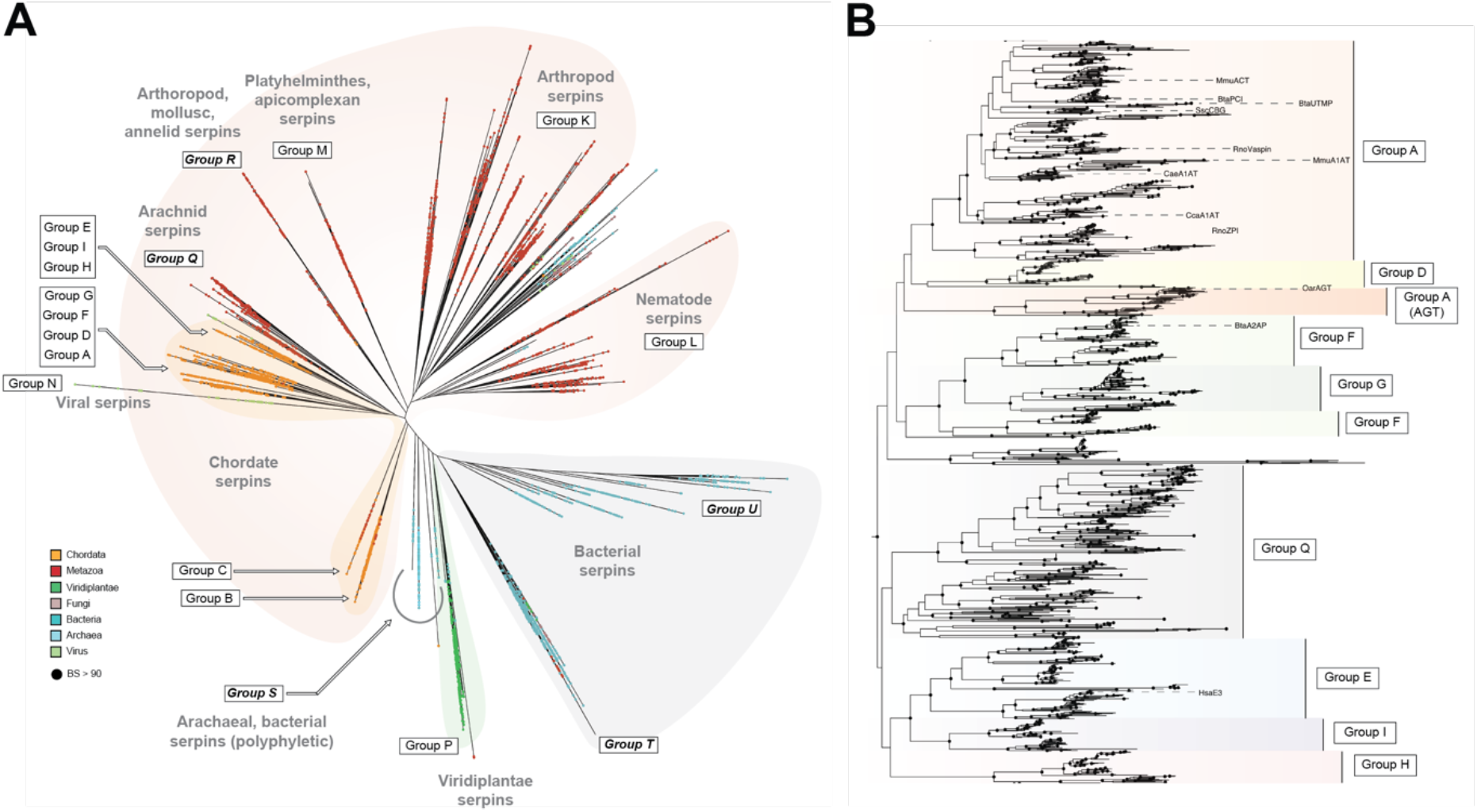
A. An unrooted maximum likelihood phylogeny of the serpin superfamily. Tips are colored by taxonomic classification and nodes with branch supports (BS) >90 are represented as solid black circles. Well characterised chordate serpin groups A, B, C, D, E, F, G, H and I, known *a priori* to belong to distinct lineages, are highlighted, as well as Groups K, L, M and O. New families defined in this study are highlighted in bold. The phylogeny is available in Newick format with full branch support values in S.I. **B**. Subtree of chordate serpin Groups A, D - I. Branch color distinguishes each Group within the subtree and functionally characterised tips are labelled.

We inferred the serpin superfamily phylogeny by maximum likelihood (ML). Tree inferences were performed with five independent replicates using the free-rate, approximated mixture model LG+R10+F+C10 that explicitly models heterogeneity in both evolutionary rates and amino acid substitution processes across sites using the posterior mean site frequency (PMSF) approximation (Wang et al. 2018). The diversity of sequences in our dataset and phylogenetic complexity of modeling evolution in the serpin superfamily warranted the use of such a complex model; indeed using approximated mixture models with PMSF profiles has been shown previously to ameliorate artifactual long branch attraction in ML phylogenies (Wang et al. 2018). Of the resulting five topologies, three were rejected by the approximately unbiased (AU) test conducted to 10000 replicates and two were statistically indifferent at explaining the alignment data (Shimodaira 2002). The single best topology that is presented in Figure 3A and used in subsequent analyses was selected by general congruence with previously published serpin phylogenies (Irving et al. 2000; Heit et al. 2013; Krem and Di Cera 2003).

The phylogeny presented in Figure 3A allowed us to define 5 previously uncategorised families of evolutionarily distinct serpin Groups (Groups Q - U) that belong to clades previously unresolved in phylogenies of the serpin superfamily. Many of the serpin families classified as single groups in previous phylogenetic studies are also much more diverse than originally credited. The Group K arthropod serpins, for example, appears to consist of at least five indepedendant major lineages. However, for simplicity, we maintain the original classification scheme for these groups as their functional diversity remains unclear and they are poorly represented by experimentally annotated serpins.

This phylogeny illustrates that the serpin superfamily can be conceptually split into three major groups: one comprising bacterial serpins, one comprising plant serpins and the other consisting of predominantly metazoan serpins. A smaller polyphyletic group of prokaryotic serpins, including Archaeal serpins (Group S), is also present. The branch that splits the metazoan serpins from the plant and bacterial serpins is the mid-point of the tree and is the most logical position for the root. Serpins belonging to Groups B, P and S form distinct clades around the root and are the most ancestral members of the superfamily. Metazoan serpins descended from a single common ancestor shared with the Group B serpins and are split into chordate (Groups A, D-I) and non-chordate lineages (Groups K-R) that diverged from one another. Most prokaryotic serpins belong to two lineages (Groups T and U) that shared a single common ancestor early in the superfamily’s history. Fungal serpins, together with some prokaryotic and non-chordate metazoan serpins are not resolved as a defined clade, but are instead orphan sequences that were not given a classification.

### Serpins in chordates

The molecular evolution of serpins in chordates has been studied extensively (Kumar and Ragg 2008; Irving et al. 2000; Heit et al. 2013; Krem and Di Cera 2003). The topology of the chordate serpin subtree (Figure 3B) is concordant with the established understanding of serpin function and evolution among higher eukaryotes. We find that Group B and C serpins are indeed ancestral to Groups A and D-I, which together form a divergent monophyletic clade. Groups A and D, F and G, and E,I and H each share a recent common ancestor; however, the cladistic inclusion of Group H with Groups E and I is only marginally supported. Groups B and C belong to sister clades that share an older common ancestor with the other chordate serpin lineages. With the exception of Group H, the topological placement and composition of each of these clades is independently supported by common patterns of intron-exon splice sites and microsynteny (Kumar and Ragg 2008) that were not considered during phylogenetic inference, demonstrating the robustness of the chordate cladistic inclusions.

As the standard for chordate serpin nomenclature is derived from cladistic grouping, the divergence of angiotensinogen (AGT) from the last common ancestor of Groups A (excluding AGT) and D (supported by BS > 99) means that current classification standards should be amended to either recognise AGT as evolutionarily distinct from Group A, or redefine Group D as redundant within Group A. AGT is not a canonical protease inhibitor and instead functions in blood plasma redox sensing and blood pressure regulation, highlighting its divergence from a canonical Group A serpin (Zhou et al. 2010; Yan et al. 2019).

### Serpins in invertebrates

Non-chordate metazoan serpins are the largest and most diverse group in the superfamily. Despite this, serpins from non-chordate metazoans are paradoxically among the most disproportionately under-represented in literature and reviewed databases.

Most characterised arthropod serpins belonging to Group K regulate innate immunity. For example, when challenged with infection, *Tenebrio molitor* serpin48 regulates the Toll pathway (Park et al. 2011) and *D. melanogaster* serpin 27A and *Aedes aegypti* serpin2 localize melanin to the site of infection *via* regulation the prophenoloxidase proteolytic cascade (An et al. 2013; An, Budd, et al. 2011). Serpin1J regulates both innate immune responses in *Manduca sexta* (An, Ragan, et al. 2011; Jiang et al. 2003), indicating that broad protease specificity and functional generality can be a feature of arthropod serpins. The Group K serpins also play a role in development. Serpins 16, 18 and 22 from *Bombyx mori* regulate in silk-gland development (Guo et al. 2015) and *Drosophila melanogaster* serpins 42 and 27A are involved in developmental protein maturation (Richer et al. 2004) and dorsal-ventral partitioning during embryonic development (Hashimoto et al. 2003; Richer et al. 2004). Many of these paradigms are shared with homologous nematode serpins belonging to Group L (Pak et al. 2004).

Non-chordate metazoan serpins have also evolved functions in the regulation of exogenous proteases that accommodate parasitic lifestyles or comprise major components in venom. Many of the characterised platyhelminthes serpins belonging to Group M inhibit endogenous and host proteases to dampen inflammation upon infection (Lopez Quezada et al. 2012), alter the proteolytic environment to optimise host penetration (Molehin et al. 2014), or to evade host immune responses (Yang et al. 2014; Ghendler et al. 1994). The inclusion of apicomplexan serpins, which remain largely uncharacterised with some exceptions (Fetterer et al. 2008), within Group M leads us to speculate that this clade comprises serpins predominantly involved in host invasion and accommodating a parasitic lifestyle; however, the functional diversity of this clade remains obscure in our analysis as relatively few Group M serpins have been characterised. Notably, some of the serpins in Group L likely perform similar functions in parasitic nematodes (Toubarro et al. 2013; Bennuru et al. 2009), and Group K serpins from the parasitoid wasp *Leptopilina boulardi* suppress innate immune responses in *Drosophila* hosts (Colinet et al. 2009), indicating that serpins involved in modulating host immunity and the parasitic microenvironment have emerged independently in multiple lineages over a broad taxonomic range of hosts. Although we maintain the original Group K nomenclature of the arthropod serpins defined by (Irving et al. 2000), here we expand the classification of the family into 5 major lineages, Groups K1 - K5 (Supplementary Figure 4).

**Figure 4.**
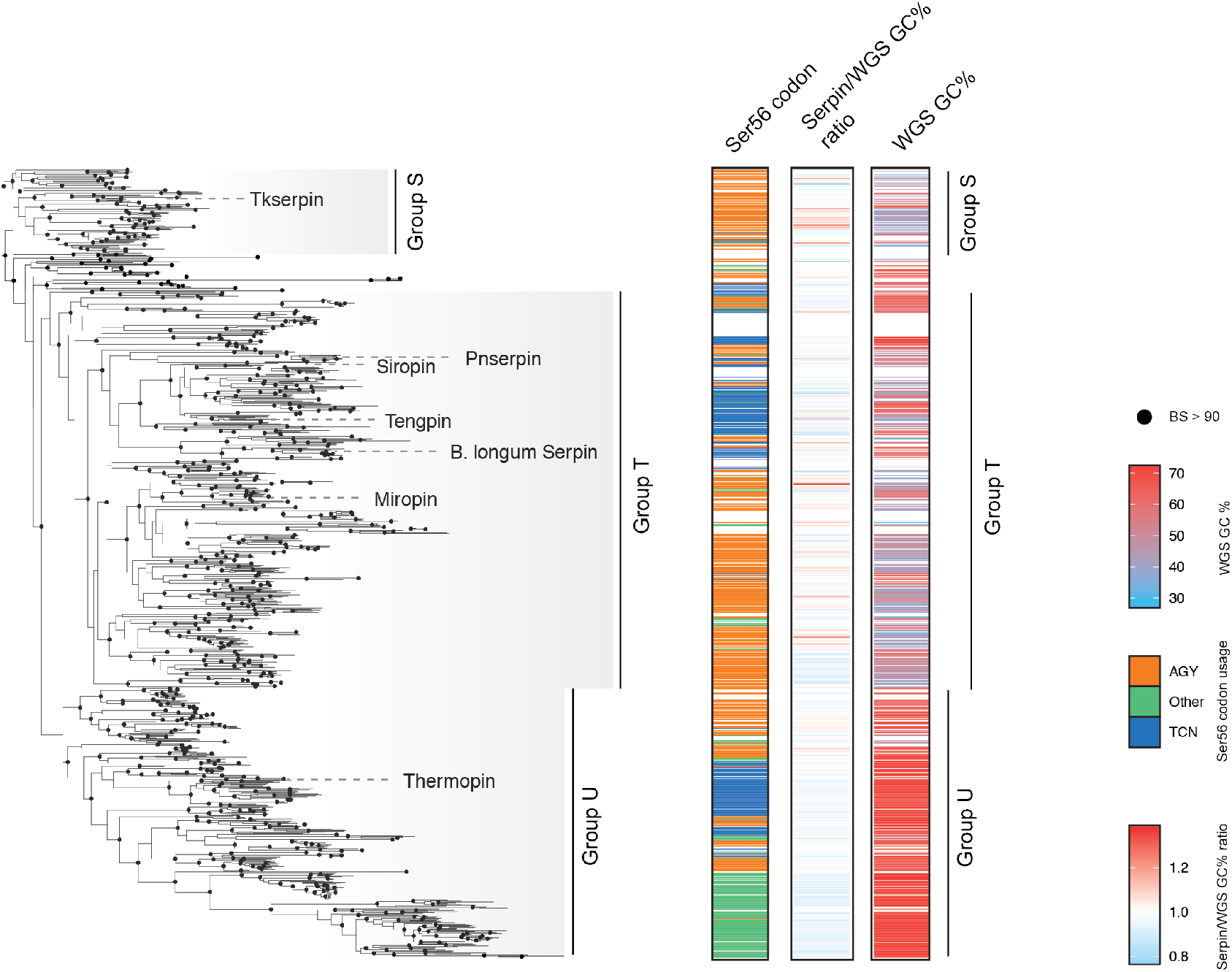
Genomic and molecular markers of horizontal gene transfer. Where available, codon usage for conserved Ser56, whole genome sequence (WGS) GC% and the ratio of serpin gene sequence GC% to WGS GC% are mapped to prokaryotic serpins belonging to Groups S-U. There are no systematic anomalous ratios in serpin gene GC% to WGS GC% over a wide range of whole genome sequence GC contents (<30% - >70%). Likewise, there is no systematic evidence that Groups S-U were acquired from chordates from the pseudo-conserved Ser56 codon usage in prokaryotes.

Among the new serpin families we define in this study are the arachnid Group Q serpins. A peculiarity among the Group Q serpins is their incongruent placement as a sister group to the chordate Groups H, I and E. A majority of serpins that make up this clade belong to Ixodid ticks, as well as few serpins from the genera *Megacormus* (scorpion), *Tityus* (scorpion), *Parasteatoda* (spider), *Stegodyphus* (spider), and *Sarcoptes* (mite, including *S. scabiei*). The Group Q serpins are also highly similar to chordate serpins and cluster together in SSNs at stringent similarity thresholds, indicating molecular homoplasy between them. It is curious that the majority of taxa that belong to Group Q are chordate parasites (ticks and mites) that have evolved serpins to inhibit host proteases in order to evade adaptive immune responses (Xu et al. 2019; Prevot et al. 2009; Chmelar et al. 2011) or prolong feeding by disrupting hemostatic proteolytic cascades (Mulenga et al. 2013; Chmelar et al. 2011). There is also evidence to suggest that serpins from Group Q form a component of venom in some free-living arachnids (scorpions, spiders) (Kazemi and Sabatier 2019; Gremski et al. 2010), although the precise function of these proteins remains unknown.

### Serpins in plants

Unlike serpins from other major taxonomic groups, all plant serpins belong to a single, well defined and homogenous clade that diverged from a single common ancestor. We maintain the original Group P nomenclature (Irving et al. 2000) that defines the plant serpins as a monophyletic family and is congruent with our cladistic groupings. Like their metazoan counterparts, plant serpins occupy diverse inhibitory and non-inhibitory functions (Cohen et al. 2019). Barley protein Z, for example, is a major storage protein during grain filling with few other putative functions (Hejgaard et al. 1985). Indeed, grain serpins can comprise as much as 4% of the total storage protein in monocot grains where they are both non-inhibitory storage proteins and protease inhibitors that protect storage proteins from proteolytic degradation by endogenous proteases (Evans and Hejgaard 1999). In the cytoplasm, inhibitory plant serpins, such as *Arabidopsis thaliana* Serpin1 (*At*serp1) inhibit ectopic cysteine proteases following vacuolar collapse from abiotic stress (Koh et al. 2016; Lampl et al. 2010) or hypersensitive response when challenged by a pathogen (Lampl et al. 2010; Lema Asqui et al. 2018), thus attenuating programmed cell-death. Plant serpins may also play a role in defence against predation by inhibiting digestive proteases in the midgut of herbivorous arthropods such as aphids (Yoo et al. 2000).

### Serpins in prokaryotes

The physiological role of serpins in many prokaryotes is enigmatic. Prokaryotes lack complex proteolytic signalling pathways and do not utilise the regulatory networks that serpins are often part of in higher eukaryotes. Despite this, serpins are found sparsely distributed throughout some bacterial phyla, with most belonging to *actinobacteria, proteobacteria, bacteroidetes* and *firmicutes*. Whereas many of the prokaryotic serpins are indistinguishable from chordate serpins in SSNs, prokaryotic serpins belong to three major families that are phylogenetically distinguished from the chordate serpins. We define these three major prokaryotic groups as S-U. Most bacterial serpins belong to Groups T and U, which shared a common ancestor early in the evolution of the superfamily. Group T is represented by miropin, siropins from *Eubacterium sirium* (Mkaouar et al. 2016), *Bifidobacterium longum* serpin (Ivanov et al. 2006) and tengpin from *Thermoanaerobacter tengcongensis* (Zhang et al. 2007). The only functionally characterised member of Group U is thermopin, belonging to *Thermobifida fusca* (Fulton et al. 2005). Sequence analysis indicates that most of the Group U and T serpins are inhibitory; however a clade of *Streptomyces* serpins belonging to Group U deviate from a canonical inhibitory serpin in conserved regions of the breach and shutter, suggesting that they may have evolved a non-inhibitory function. Despite their highly diverged sequences, these *Streptomyces* serpins share significant homology (E-value <10e-10) with other prokaryotic and eukaryotic serpins in an NCBI BLAST search and many retain the consensus RCL-hinge motif expected in an inhibitory serpin (Hopkins et al. 1993; Irving et al. 2000).

Some bacterial serpins appear to play a role in host-commensal or host-pathogen interaction. Miropin, for example, has recently been demonstrated to efficiently inhibit human plasmin, thereby protecting invading *T. forsythia* cells from plasmin mediated degradation and attenuating fibrinolysis in the pathogen’s local environment (Sochaj-Gregorczyk et al. 2020). Siropins belonging to the commensal bacterium *E. sireum* have been shown to inhibit two human proteases, human neutrophil elastase (HNE) and proteinase3, that are abundant in gastrointestinal (GI) tract where *E. sireum* is prevalent (Mkaouar et al. 2016) and *Bifidobacterium longum* and *B. breve*, both commensal members of the GI microbiome, express serpins capable in inhibiting mammalian proteases present in their microenvironment (Turroni et al. 2010; Ivanov et al. 2006). To investigate the prevalence of prokaryotic serpins in different human microbiota, we performed chemically-guided functional profiling (CGFP) (Kaminski et al. 2015) on MCL clusters of prokaryotic serpin sequences. The CGFP workflow matches sequence markers from SSN clusters to markers from metagenomic reads of different human microbiome samples, thus detecting sequence similarity group presence among human microbiota. Many prokaryotic serpins beyond those that have been experimentally characterised have a significant presence in the human oral and GI microbiomes (Supplementary Figure 5). The most abundant of these belong to Group T, which includes serpins from many known pathogens and commensal bacteria (such as *Prevotella spp*., *Corynebacterium spp*., and *Lachnospiraceae spp*.) and are closely related to miropin, *Bifidobacterium spp*. serpins and siropins. Many serpins belonging to Group U also have either a presence, or close homologues with a presence, in various human microbiomes. No serpins belonging to Group S, which is composed of both archaeal and bacterial serpins, were detected with significance, indicating that they are likely functionally distinct from the Group S and T serpins.

**Figure 5.**
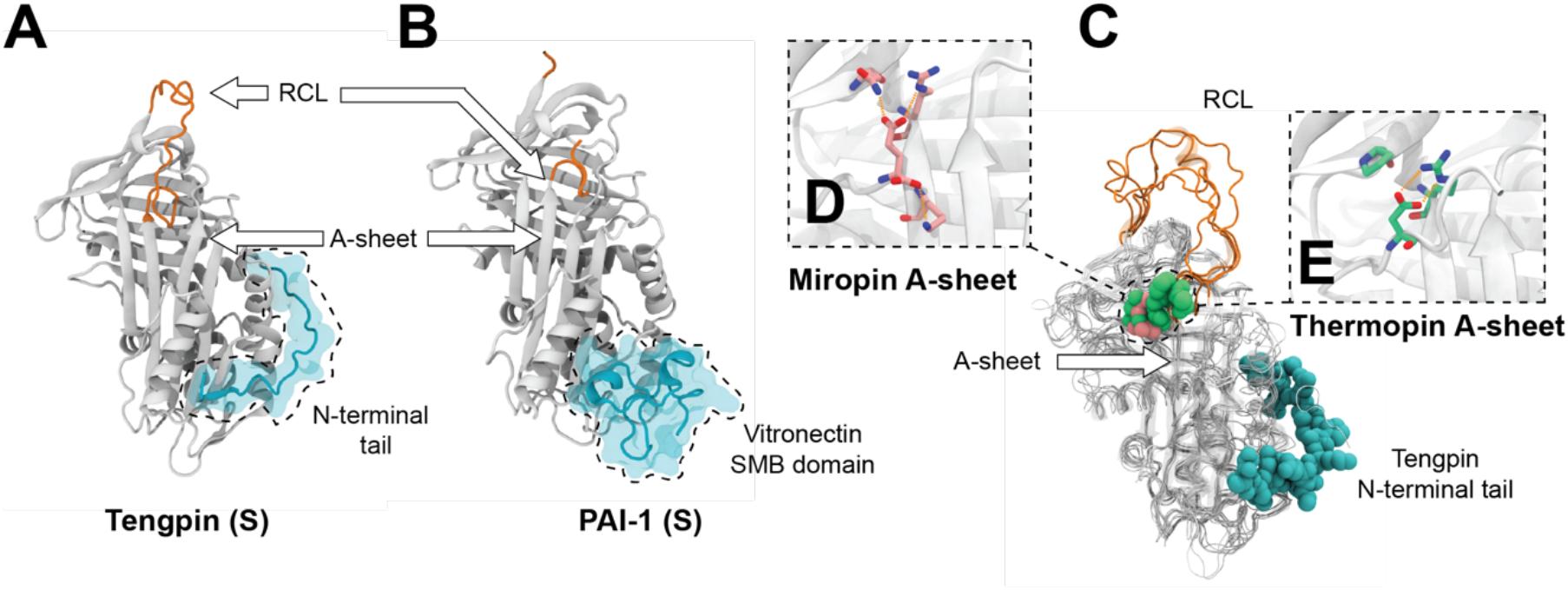
Structural evidence of convergent evolution in the serpin superfamily. **A**. Cartoon representation of tengpin in the stressed native state. The N-terminal extension and its contacts in the A-sheet are highlighted, among other structurally important positions, such as the A-sheet and RCL (PDB: 2PEE). **B**. Cartoon representation of PAI-1 in the stressed native conformation, stabilized by the SMB domain of vitronectin (highlighted in cyan) (PDB: 1OC0). **C**. A structural alignment of tengpin (PDB: 2PEE), miropin (PDB: 5NCS), thermopin (PDB: 1SNG), SCCA-1 (PDB:2ZV6) and AT3 (PDB:3KCG) highlighting the convergent molecular mechanisms exploited to stabilize the native conformation. **D**,**E**. View of analogous interactions in the C-terminal tails of miropin and thermopin that stabilize the breach, respectively.

### Serpins are an ancient superfamily

The common ancestry shared between prokaryotic serpins from Groups T and U suggests that bacterial serpins descended from a single common ancestor that was distinct from the last common ancestor of the chordate serpins. We found no phylogenetic evidence supporting the hypothesis that prokaryotes acquired serpins *via* HGT as no phylogenetic topology (including those that were rejected by the AU-test) placed prokaryotic serpins within the chordate serpin lineages, despite their high similarity and co-clustering in SSN.

Independent of phylogenetic topology, there are often other molecular markers that may indicate that horizontal gene transfer has occurred. We found few and non-systematic anomalous GC%_gene_/GC%_genome_ ratios (>1.2 or <0.8) within bacterial serpin genes (Supplementary Figure 6). This included many species of *Streptomyces* and *Mycoplasma* that are characterised by high (>70%) and low (<30%) genomic GC content respectively, suggesting that either horizontal gene transfer has not occurred or that genetic transfer events were ancient enough to have ameliorated in extant DNA.The protostome-deuterostome codon usage dichotomy at the conserved position Ser56 (Krem and Di Cera 2003) also provided no evidence of horizontal gene transfer. Ser56 is only marginally conserved in prokaryotic serpins and no particular lineage strictly adheres to either the archaic protostome ACY codon or the TCN codon that is fixed in chordates. Instead, bacterial lineages that use either codon are generally separated by clades where Ser56 has been lost, parsimoniously demonstrating the loss of Ser56 under one codon before re-fixation under the other. This, corroborated by the absence of a conceivable mechanism of genetic transfer between a chordate and many of the prokaryotes that feature serpins (such as free-living, environmental extremophiles) and the lack of phylogenetic evidence supporting the HGT hypothesis, indicates that the serpin superfamily is ancient and emerged in the last common ancestor of the major bacterial lineages at the latest, independently of eukaryotes. This conclusion is equally supported by the alternative phylogenetic topology that failed rejection by the AU-test (Supplementary Figure 7).

### Structural analysis of the serpin superfamily reveals evidence of convergent evolution

A structural comparison between bacterial and chordate serpins reveal several distinct structural adaptations to selective pressures, some of which are shared between families. The majority of these are solutions to the unique structure-function challenge faced by serpins; specifically, how to maintain the relative stabilities of the native and cleaved states in order to function as inhibitors. Serpins from thermophilic bacteria have the added complexity of maintaining this balance at high temperatures. Thermopin, from the moderate thermophilic bacteria *T. fusca* (optimum growth temperature 55 °C), contains a unique C-terminal extension that packs against the face of the top of the A-sheet (Figure 5A, (Fulton et al. 2005)). This tail interacts with highly conserved residues in the breach that are known to be critical for controlling serpin stability and function. Tengpin from the extremophilic prokaryote, *T. tengcongensis* (optimum growth temperature 75 °C), contains an N-terminal tail that binds to a hydrophobic patch near strand s1A on the body of the protein, stabilising the native state (Zhang et al. 2007). Indeed, evolution has overcome the thermodynamic challenges imposed by the serpin fold and function many times. Aeropin, from the hyperextremophile archaea *Pyrobaculum aerophilum* (growth temperature 100 °C) (Cabrita et al. 2007), features a C-terminal tail of similar length to that of thermopin and two disulfide bonds that are structurally non-homologous to disulfides in mesophilic serpins, such as the single disulfide bond in miropin. In both cases, the disulfides contribute significant stabilisation energy to the native conformation. Interestingly, structural comparison also shows that the C-terminal region of miropin forms a contact with the breach through a glutamic acid residue, analogous to the aspartate breach interaction in the non-homologous C-terminal tail of thermopin (Figure 5C,D). The phylogenetic distance between thermopin (Group U) and the Group T thermophilic serpins from *Pyrobaculum spp*. and tengpin and the analogy between aeropin and miropin disulfide bonds would suggest that thermophilic serpins from different evolutionary backgrounds have independently acquired different and often convergent molecular mechanisms of accommodating the thermodynamic balance required for serpin metastability.

The biophysical solutions that accommodate metastability have emerged elsewhere in serpin function, as well. The structurally-sensitive region of the serpin fold that is stabilized by the N-terminal tail of tengpin is the same site on the mammalian serpin PAI-1 bound by the somatomedin B (SMB) domain of the plasma protein vitronectin (Zhou et al. 2003; Zhang et al. 2007); Figure 5B), which functions to stabilise the metastable native state of PAI-1 and thus regulate its function (Declerck et al. 1988). The structural analogy between the N-terminal tail of tengpin and the SMB binding site of PAI-1 is a stark example of structural convergence where the same biophysical defect in the serpin fold is overcome by analogous mechanisms to achieve different biological results (thermostability in tengpin and regulation by vitronectin in PAI-1). This structural analogy may also suggest an interesting potential mechanism for the evolution of metastability and serpin-cofactor interactions. We can speculate that tengpin represents an ancestral origin of the conformational control mechanism observed in mammalian serpins such as PAI-1. In such a hypothesis, the progenitor to PAI-1 consisted of an N-terminal vitronectin domain that has since been removed during its evolution. Such genetic separation of the N-terminal tail from the serpin domain could transfer control of metastability to a separate protein cofactors (Zhang et al. 2007), permitting more biologically complex mechanisms of regulation.

Perhaps the most compelling evidence supporting the HGT hypothesis is the apparent structural similarity between the solved crystal structures of miropin and the chordate serpin AT3 (Goulas et al. 2017). Indeed, structural alignment between miropin in a native conformation and tengpin, thermopin, AT3 and SCCA-1 reveals that miropin shares the lowest RMSD with AT3 (1.3 Å) and SCCA-1 (1.6 Å) (compared to 1.8 Å with thermopin and 1.9 Å with tengpin). This is only true when using a single structural model of AT3 in a ternary michaelis complex with thrombin and heparin (PDB: 3KCG). When a more complete conformational ensemble of AT3 is considered in structural analysis with native miropin, including monomeric apo-AT3 structures (PDBs: 2ANT, 1TB6, 1T1F), RMSDs range from 1.6 Å - 4.1 Å and are on average 2.7 Å. The perceived close similarity between miropin and AT3 is likely an artifact stemming from the structural comparison of biologically distinct conformations of miropin (monomeric native) and AT3 (ternary michaelis complex). It is nonetheless an interesting observation (Goulas et al. 2017) that miropin does appear to share more structural features with chordate serpins than what may be expected from proteins that share often <30% sequence identity. In the absence of phylogenetic or molecular evidence supporting the HGT hypothesis, it is most likely that this is yet another example of structural convergent evolution within the serpin superfamily. We hypothesize that structural convergence is such a persistent evolutionary phenomenon among serpins because of the inherent thermodynamic balance that must be maintained by the serpin fold for metastability. It is conceivable that there are only few biophysical solutions to a structure where both stressed and relaxed conformations are accessible. Divergence from that structure would generate non-inhibitory serpins, resulting in a fitness landscape with clearly defined fitness peaks that have been converged independently over the evolutionary history of the serpins.

### The evolution of serpin function and location

By identifying N-terminal signal peptides, we were able to classify serpin sequences as likely intracellular, extracellular, periplasmic (bacterial serpins only) or membrane anchored (prokaryotic serpins only) (Supplementary Figure 8). The ancestral-like eukaryotic serpins are dominated by sequences that lack N-terminal signal peptides and likely function intracellularly, including the Group P plant serpins and the chordate Group B serpins. Extracellular serpin expression emerged within the diverged metazoan lineages, including chordate Groups A, D-H and Groups K-L of the arthropod and non-chordate metazoan lineages. Notably, many of the Group L nematode serpins lack signal peptides and appear to function in the cytosol. The lower degree of biological organization among prokaryotes makes the functional distinction between intracellular and extracellular prokaryotic serpins less significant than that of eukaryotic serpins. Most prokaryotic serpins feature N-terminal transport peptides, although we were unable to identify systematic trends that relate phylogenetic placement to cellular localisation and function among prokaryotic serpins.

We hypothesize that ancestral serpins were intracellularly localised and occupied biologically rudimentary functions, such as protecting cells from promiscuous proteolysis. Intracellular localisation is likely a trait that was retained in the ancestral-like extant eukaryotic serpins, which diverged from ancestral functions as biological systems and cellular biology became more complex over evolutionary history. Indeed, there are functional similarities between extant Group B serpins, many of which protect cells from ectopic endolysosomal proteases and a hypothetical primordial serpin that serves solely cytoprotective functions. This paradigm also extends to other ancestral-like Group P plant serpins. The *A. thaliana At*serp1, for example, inhibits proteases that have escaped from ruptured vacuoles and ER bodies, thereby protecting cells from programmed death (Lampl et al. 2010). In contrast, the extracellular eukaryotic serpins (as well as few intracellular serpins that do not belong to the ancestral-like lineages, such as HSP47) have diverged into functions that accommodate higher biological organization, such as hormone transport, hemostasis and immunity, among others. The antithrombin Group C lineages occupy an important position in chordate serpin evolution. Unlike other chordate serpins, those belonging to Group C are not unanimously intracellular or secreted and topologically belong to a monophyletic clade that separates the last common ancestors of Group B and Groups A, D-I, indicating that antithrombin-like serpins were the first to appear extracellularly and likely bridge intracellular and extracellular serpin functions.

Such conclusions are more challenging to draw regarding prokaryotic serpins. Due to their scarce presence in reviewed databases, the extent of functional diversity within Groups S-U is obscure. We can hypothesize, however, that many of the bacterial serpins belonging to Groups T and U have diverged functions involved in host-microbe interaction in vertebrate microbiota, whereas the ancestral-like Group S serpins are undetectable in microbiome metagenomes (Supplementary Figure 5). According to our proposed model of serpin evolution, the Group S ancestral-like serpins are likely functional housekeepers that protect the cell from either endogenous or exogenous environmental proteases, although this conclusion will remain uncertain until our understanding of prokaryotic serpins advances. Additionally, it remains perplexing why most prokaryotic lineages have lost serpins over evolutionary history. The lack of commonalities in biochemical niche and life-history traits among the few prokaryotic lineages that have retained serpins makes this difficult to speculate on; such questions should be the focus of future studies on the evolution of prokaryotic serpins.

## Conclusion

In this work we have leveraged the substantial structural characterization of members of the serpin superfamily to allow us to create a comprehensive phylogeny. This analysis has identified a number of currently uncharacterized, or orphan-function, clades, particularly within non-chordate phlyla. In addition to these diverse and uncharacterized clades, we also observe a large central hub of serpin sequences that includes serpins from bacteria, archaea, metazoa and plants. We hypothesize that the hub-like prokaryotic serpins occupy primitive, intracellular housekeeping roles, such as controlling rampant cytoplasmic proteolysis and likely resemble ancestral inhibitory serpins. Our analysis indicates that these closely related hub proteins are therefore ancient and are similar because of convergent evolution, rather than the alternative hypothesis of HGT, for which we find little evidence. We hope that this analysis will provide new directions for research in the field of serpin biochemistry, particularly in the characterization of non-chordate metazoa and bacterial serpins.

## Methods

### Dataset collection and SSN

All 18233 amino acid sequences that comprised the serpin superfamily at the commencement of this study were retrieved from the pfam database (PF00079) using the enzyme function initiative enzyme similarity tool (EFI-EST) server (Gerlt et al. 2015; Zallot et al. 2018). An arbitrary sequence redundancy threshold of 75% sequence identity was imposed on this dataset, reducing its size to 10123 unique serpin sequences before pairwise sequence identities were computed. The resulting SSN was visualized in cytoscape (3.6) using the yFile organic force-directed network layout. Homogenous sequence clusters were defined using the Markov clustering algorithm from the clustermaker cytoscape package (Morris et al. 2011) and validated with experimental evidence from the SwissProt database. Network topology tests, such as centrality metrics, were computed within cytoscape. DNA sequences for serpin encoding genes and prokaryotic genomes were retrieved from NCBI using reference uniprot IDs and organism IDs, respectively. N-terminal signal peptides were identified using the SignalP4.0 algorithm (Petersen et al. 2011).

### Sequence alignment

All sequences that made up the SSN were retrieved from the UniProt database and inspected to confirm their classification as canonical serpins on the basis of the hinge, breach and shutter motifs. As pairwise identities between sequences in the dataset were often <15%, progressive and consistency-based alignment algorithms typically failed to produce robust multiple sequence alignments, which were benchmarked against structural alignments of representative serpin crystal structures. Instead, sequences were aligned according to ensembles of HMMs that were trained iteratively with increasing diversity of sequence data, starting initially from an HMM trained on the distribution of amino-acids tolerated at each site of the conserved serpin fold. 39 representative serpin X-ray crystal structures in the stressed conformation (PDBs: 2WXY, 2CEO, 2PEE, 3B9F, 4IF8, 2ZV6, 1JMJ, 2HI9, 1OVA, 1A7C, 3F1S, 4AU2, 5M3Y, 4GA7, 1IMV, 1SEK, 2H4R, 4DTE, 5HGC, 2V95, 1WZ9, 4AJT, 4DY0, 5C98, 1BY7, 2R9Y, 1QMN, 3STO, 3F5N, 1SNG, 1YXA, 5NCS, 3OZQ, 3LE2, 3KCG, 4R9I, 5DU3, 3PZF, 2WXW) belonging to phylogenetically diverse taxa were selected from the serpin SSN and structurally aligned with MUSTANG (Konagurthu et al. 2006; Konagurthu et al. 2010). A global HMM was built from the translation of this structural alignment into a multiple sequence alignment using the default parameters of HMMBuild from the HMMer package (Finn et al. 2011). All independent sequence clusters from the SSN (at a similarity threshold of 52.5% sequence identity) were aligned internally against the structure-guided global HMM and cluster-specific HMMs (single HMM per sequence cluster in the SSN) were built using HMMBuild. The most consensus-like sequence of each cluster was identified by scoring all sequences in that cluster against its cluster-specific HMM. The resulting 750 representative sequences (from 750 sequence clusters) were retrieved and aligned against the structure-guided HMM using HMMalign in HMMer. A guide-tree was inferred from this representative alignment in IQTREE (L.-T. Nguyen et al. 2015), using LG+R10 as the ML model (selected by ModelFinder) (Kalyaanamoorthy et al. 2017) and default parameters of the tree-search algorithm, for aligning all serpin sequences in the dataset against the representative seed alignment in UPP from the SEPP package (N.-P.D. Nguyen et al. 2015). Columns that contained residues not-resolved in crystal structures (such as the RCL and non-conserved insertions) were not aligned as part of this workflow and hence deleted from the alignment. After manual editing and deletion of poorly aligned sequences, filtering of sequences that were greater than 25% shorter or longer than the median sequence length and manual refinement, the alignment consisted of 6000 unique serpin sequences.

### Phylogenetic inference

The serpin superfamily phylogeny was inferred in IQTREE v1.61 (L.-T. Nguyen et al. 2015) and all phylogenetic inferences were performed with five independent replicates. Initial guide-trees were inferred on the final serpin alignment using default parameters (perturbation strength for nearest neighbor interchange (NNI) of 0.5, maximum 1000 iterations of tree-search, tree-search concluded after 100 unsuccessful iterations) with the sequence evolution model LG+F+R10. Each of the resulting trees (irrespective of topology) was used as a guide-tree to approximate PMSF profiles using the LG+F+R10+C10 complex model (Wang et al. 2018) with extended tree-search parameters (perturbation strength for NNI of 0.2, maximum 2000 iterations of tree-search, tree-search concluded after 200 unsuccessful iterations). Branch supports were measured by ultrafast bootstrapping approximated to either 10000 or 1000 replicates (Hoang et al. 2018). Topologies that were poorly fitted to the alignment data were rejected using the AU-test (Shimodaira 2002), which was conducted to 10000 replicates and the single best topology not rejected by the AU-test was selected according to *a priori* criteria on serpin and prokaryotic evolution (Supplementary Figure 7). All tree visualization and analysis was performed in R using the libraries ggtree (Yu et al. 2017), ape (Paradis et al. 2004), tidytree, and phylobase.

