## Supplementary Information for "A comprehensive phylogenetic analysis of the serpin superfamily"

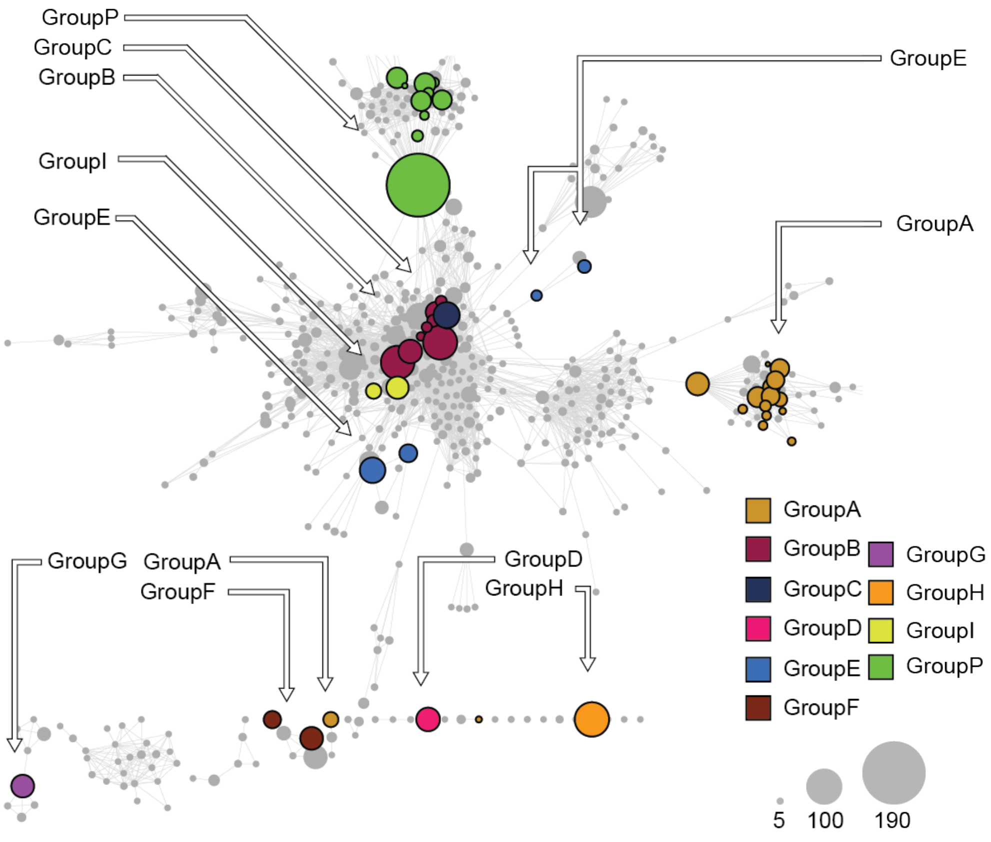


**Supplementary Figure 1. Serpin superfamily SSN with Markov clustered chordate and plant serpin families**. Each node in the SSN represents a Markov clustered group of sequences from the non-redundant full serpin superfamily SSN (main text Figure 2). Annotations were assigned to clusters of sequences according to members of that cluster with reviewed functional annotations. Serpin families are distinguished by color and node size represents the number of sequences belonging to that cluster. Serpin paralogues within a family are generally resolved as different nodes within that family.

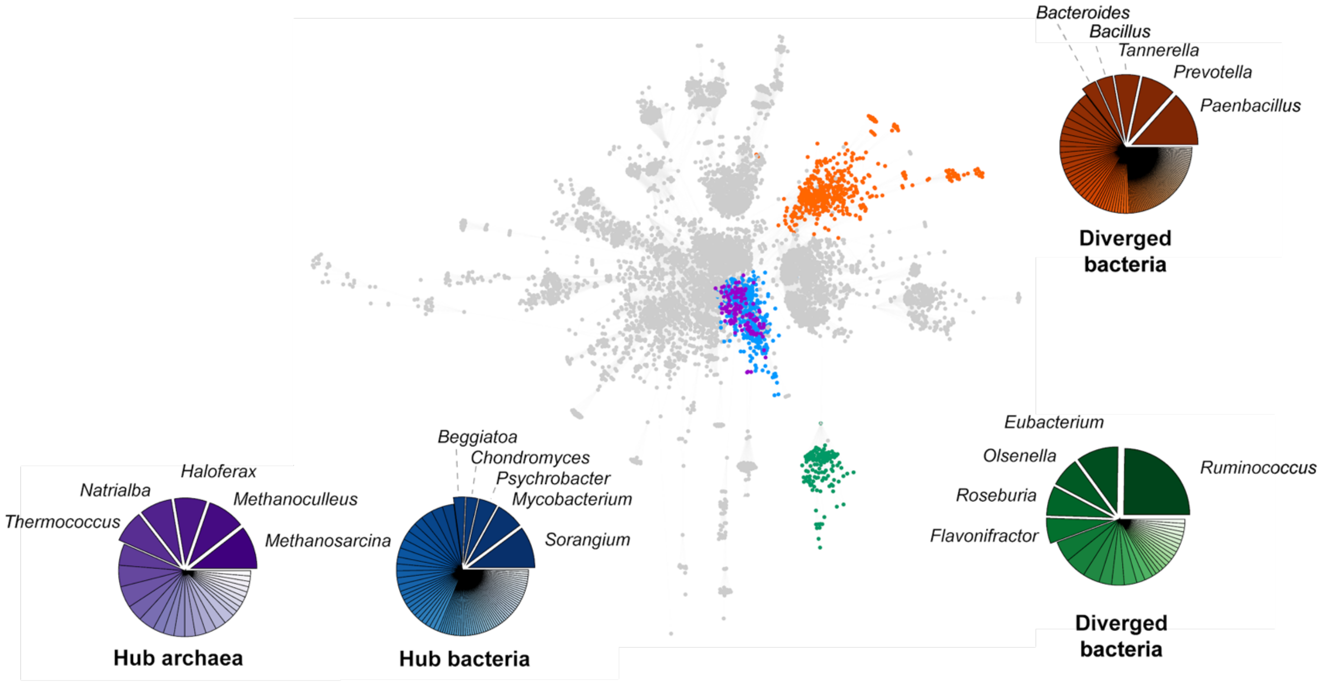


**Supplementary Figure 2. Non-redundant serpin superfamily SSN with different prokaryotic taxa**. Diverged bacterial groups (orange, green) and hub prokaryotic groups (purple, blue representing archaeal and bacterial serpins, respectively) are highlighted. Pie graphs show the taxonomic genera that serpins from each group belong to, with the most abundant labelled. Hub archaeal and bacterial serpins, which share the greatest sequence similarity with chordate serpins belong predominantly to free-living prokaryotes, whereas the diverged groups that share more distant similarity with chordate serpins belong to known pathogenic and commensal bacteria.

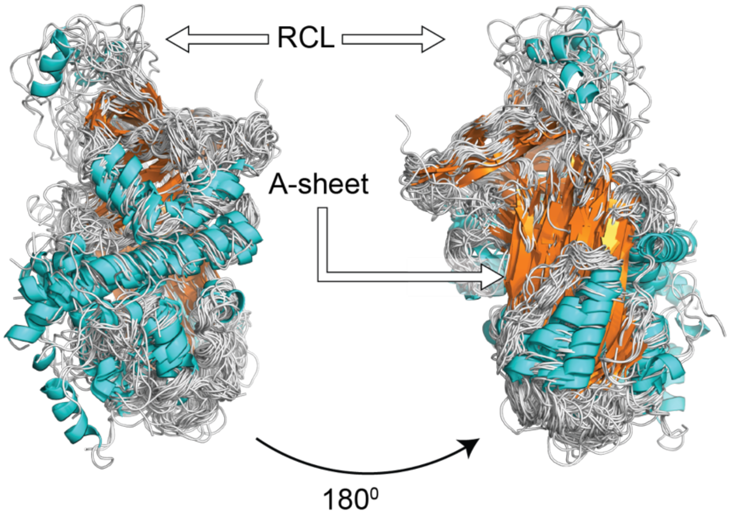


**Supplementary Figure 3. Structural alignment of 39 phylogenetically diverse serpin crystal structures used to guide sequence alignment**. Helices and sheets are represented in cyan and orange, respectively. The RCL and A-sheet are labelled. Despite extensive sequence divergence, the consensus serpin structure (excluding the RCL and non-conserved insertions) is highly conserved in the serpin superfamily.

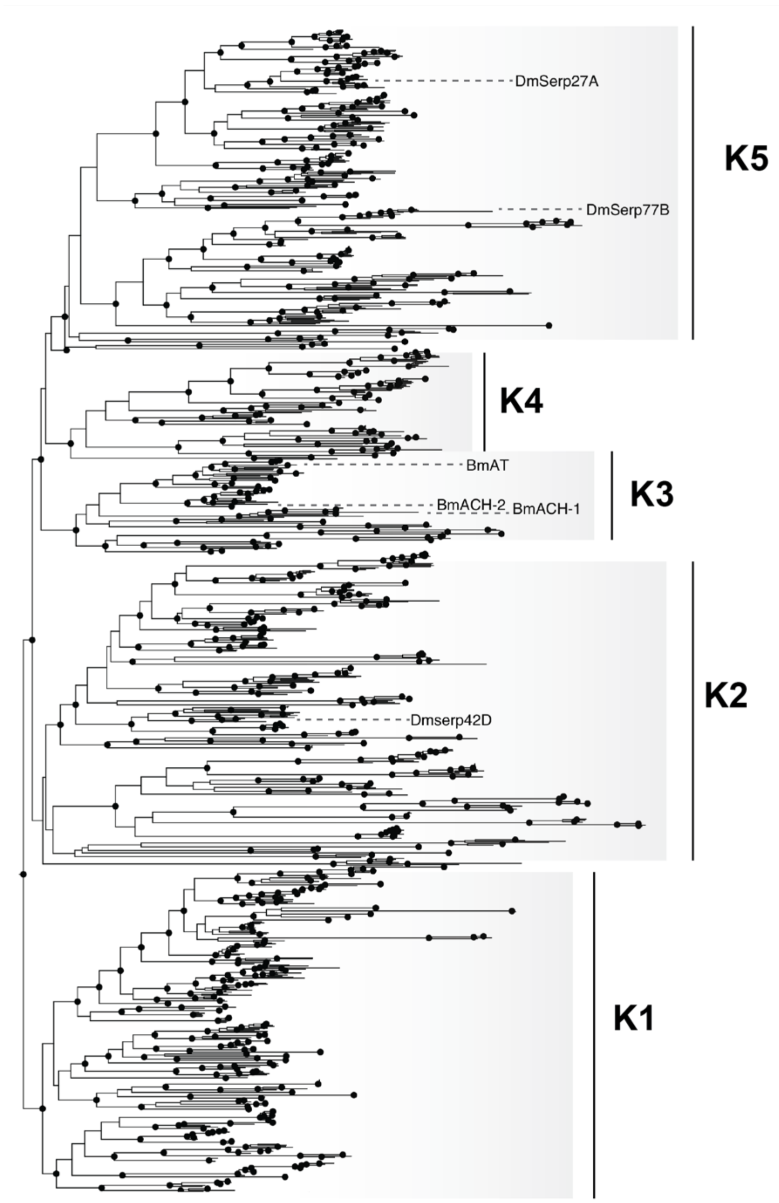


**Supplementary Figure 4. Group K subtree and family expansion** [Abbreviations Dmserp27A: *Drosophila melanogaster* Serpin 27A; Dmserp77B: *D. melanogaster* serpin 77B; BmAT: *Bombyx mori* Anti-trypsin; BmACH-2: *B. mori* Anti-chymotrypsin-2; BmACH-1; *B. mori* Anti-chymotrypsin-1; Dmserp42D: *D. melanogaster* Serpin 42D]. Solid black circles represent nodes with a BS > 90. Notable tips are labelled. Here, we expand the previously designated Group K arthropod serpin family into Groups K1 - K5, corresponding to the five monophyletic clades within the Group K family. Group K2 is represented by Dmserp42D, Group K3 is represented by *B. mori* serpins AT, ACH-1 and ACH-2, Group K5 is represented by *D. melanogaster* serpins 77B and 27A. Groups K1 and K4 lack functional representation. A group of orphan sequences belonging to fungi, protists and some bacteria has been collapsed for clarity.

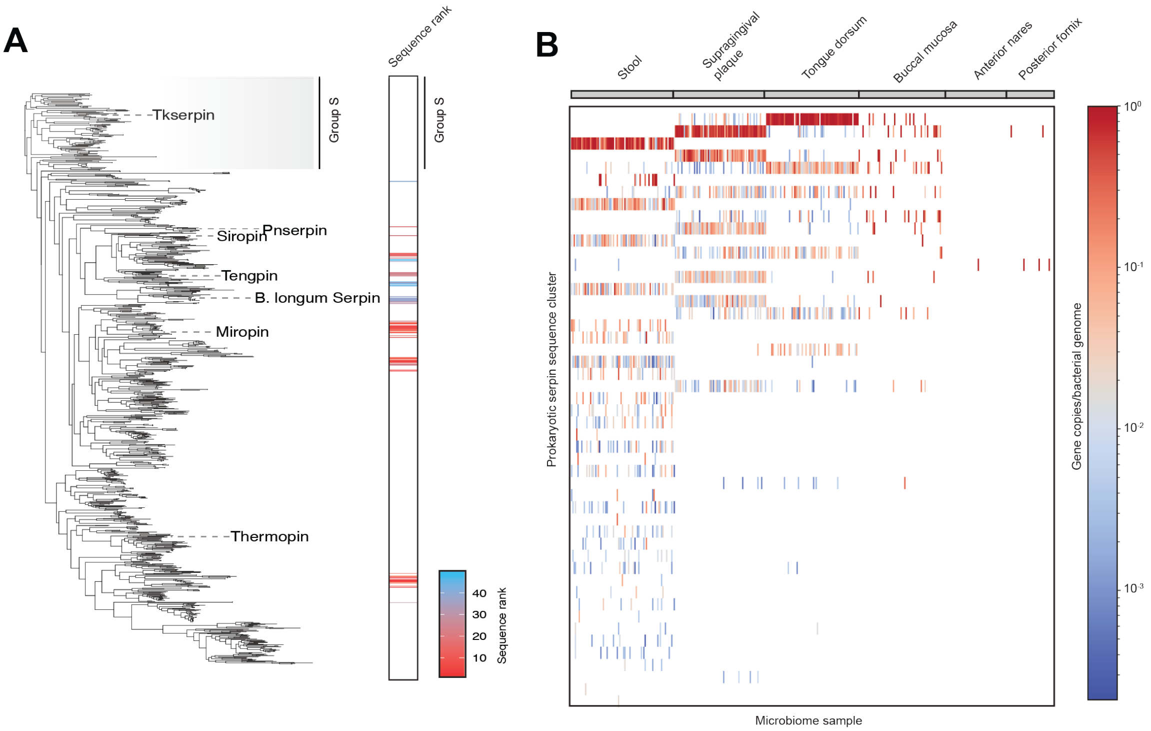


**Supplementary Figure 5. Chemically-guided functional profiling of prokaryotic serpins in different human microbiome metagenomic samples. A**. Prokaryotic serpin subtree of Groups S-U. Tips with a significant presence in different human microbiome samples are highlighted by their ranking in total gene abundance. **B**. CGFP genetic abundances in different prokaryotic serpin sequence clusters. Sequence clusters (y-axis) are ordered from the most to least abundant and colored by gene copies per microbiome genome reads (log-scale). Metagenomes (x-axis) are ordered according to the human microbiome they were sampled from. Prokaryotic serpins have a significant presence (often $10^0$ - $10^{-1}$ gene copies/genome, deep red) in the human oral and gastrointestinal microbiomes.

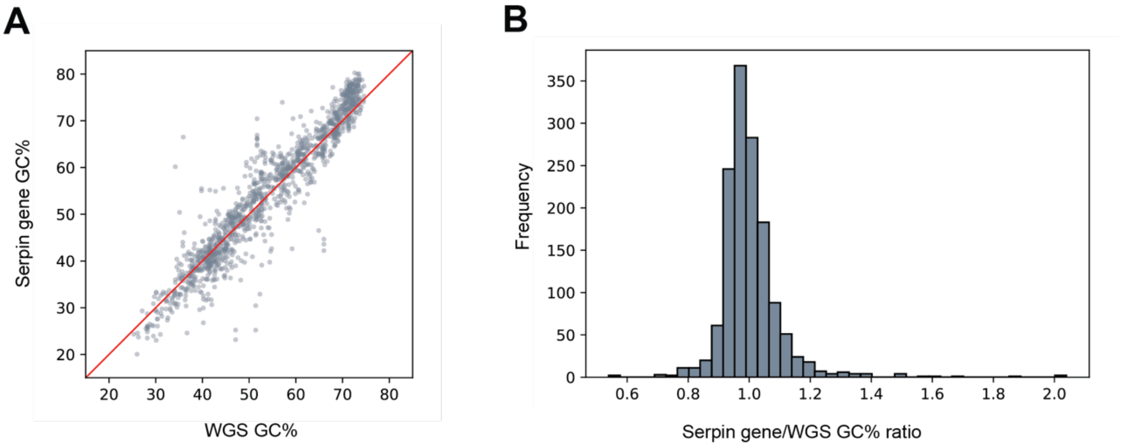


**Supplementary Figure 6. GC content distributions for whole genome sequence (WGS) and serpin encoding genes. A.** Serpin gene GC% plotted against WGS GC%. The GC content of prokaryotic serpin genes are highly correlated against the WGS GC% over a large range of WGS GC content (<30% - >70%). **B**. Histogram of serpin gene GC% to WGS GC% ratios. The distribution is normal with a mean of 1.001 and a 95% confidence interval of 0.9964 - 1.0070. Approximately 95% of observations are in the range of 0.7997 - 1.2037.

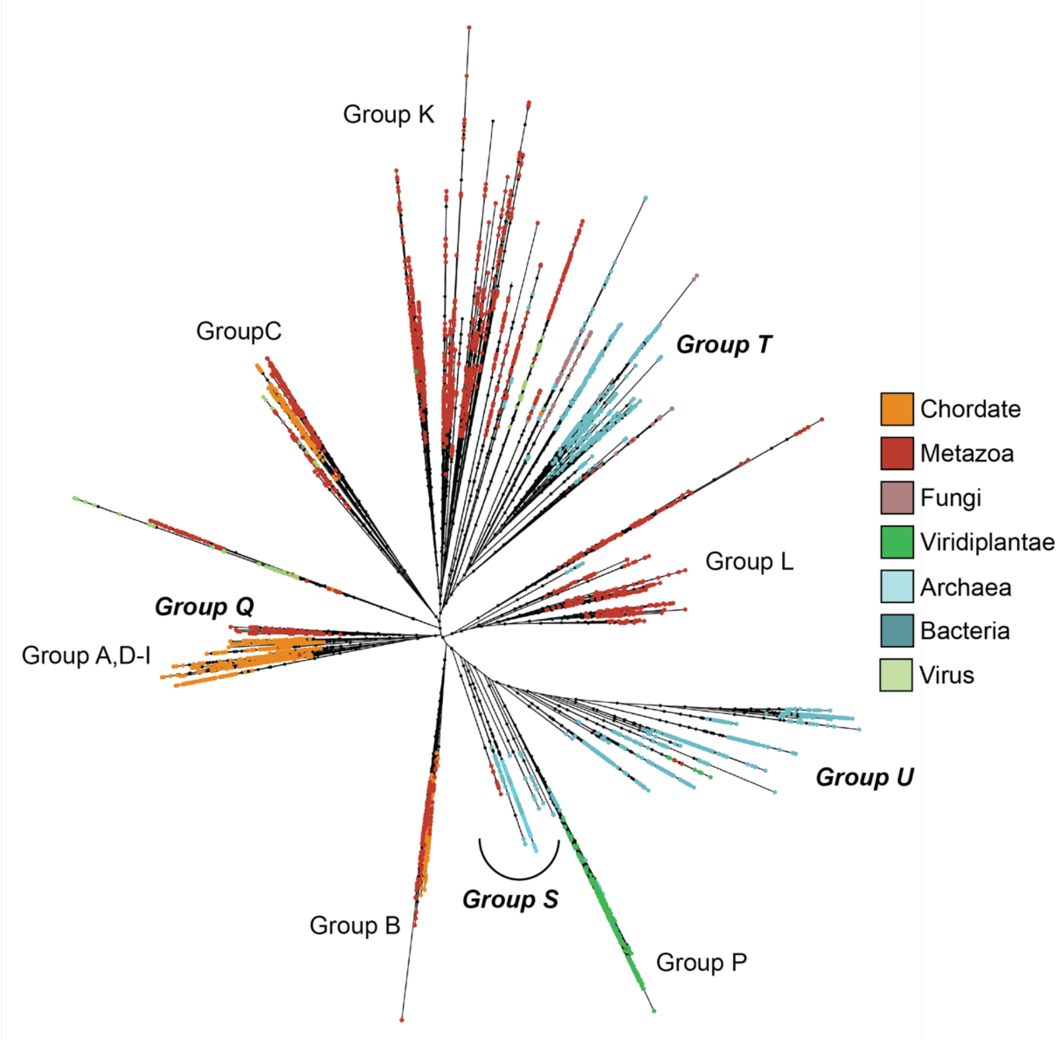


**Supplementary Figure 7**. **Alternative phylogenetic topology that passed the AU-test.** Tree-search parameters used in phylogenetic inference are the same as that presented in the main-text. The alternative topology of the serpin phylogeny that also passed the AU test similarly placed bacterial serpins as the two major, albeit polyphyletic, clades T and U that were both phylogenetically distinct from chordate serpins. Under this topology, bacterial serpins from predominantly *Actinobacteria* (clade T) appear to have evolved in parallel with and independently of serpins from *Firmicutes* and *Bacteroidetes,* among other bacterial phyla (clade U), placing the emergence of serpins in the last common ancestor of either *Actinobacteria* (diverged approximately 1 - 2 Gya), or *Firmicutes* and *Bacteroidetes* (diverged approximately 3 Gya). The polyphyletic nature of bacterial serpins under this topology is paradoxical. *Bacteroidetes* and *Firmicutes* are distantly related within the prokaryotic tree of life and the most recent ancestor shared between them was likely also the most recent bacterial common ancestor. Even in the event of horizontal gene transfer from a primordial *firmicute* to a *bacrteroidete* within clade U, the distinct phylogenetic distance between clade T from serpins in *actinobacteria* (clade U), which appear to have diverged predominantly by vertical evolution, challenges the placement of the serpin progenitor in bacteria. If clades T and U are indeed polyphyletic, how then could serpins have emerged independently in *actinobacteria* and descended vertically within that lineage billions of years after the last common ancestor of *actinobacteria* had emerged? This is one of the criteria that led us to reject this topology in favor of that presented in Figure 2, yet still consider it a statistically robust (if evolutionarily unreasonable) outcome that conservatively places the emergence of serpins in the last common ancestor of *Actinobacteria,* at the latest. Regardless, we found no evidence supporting the hypothesis of chordate-to-prokaryote xenogenous gene transfer in any of the independent inferences as no single topology placed bacterial serpins among the chordate serpin lineages.

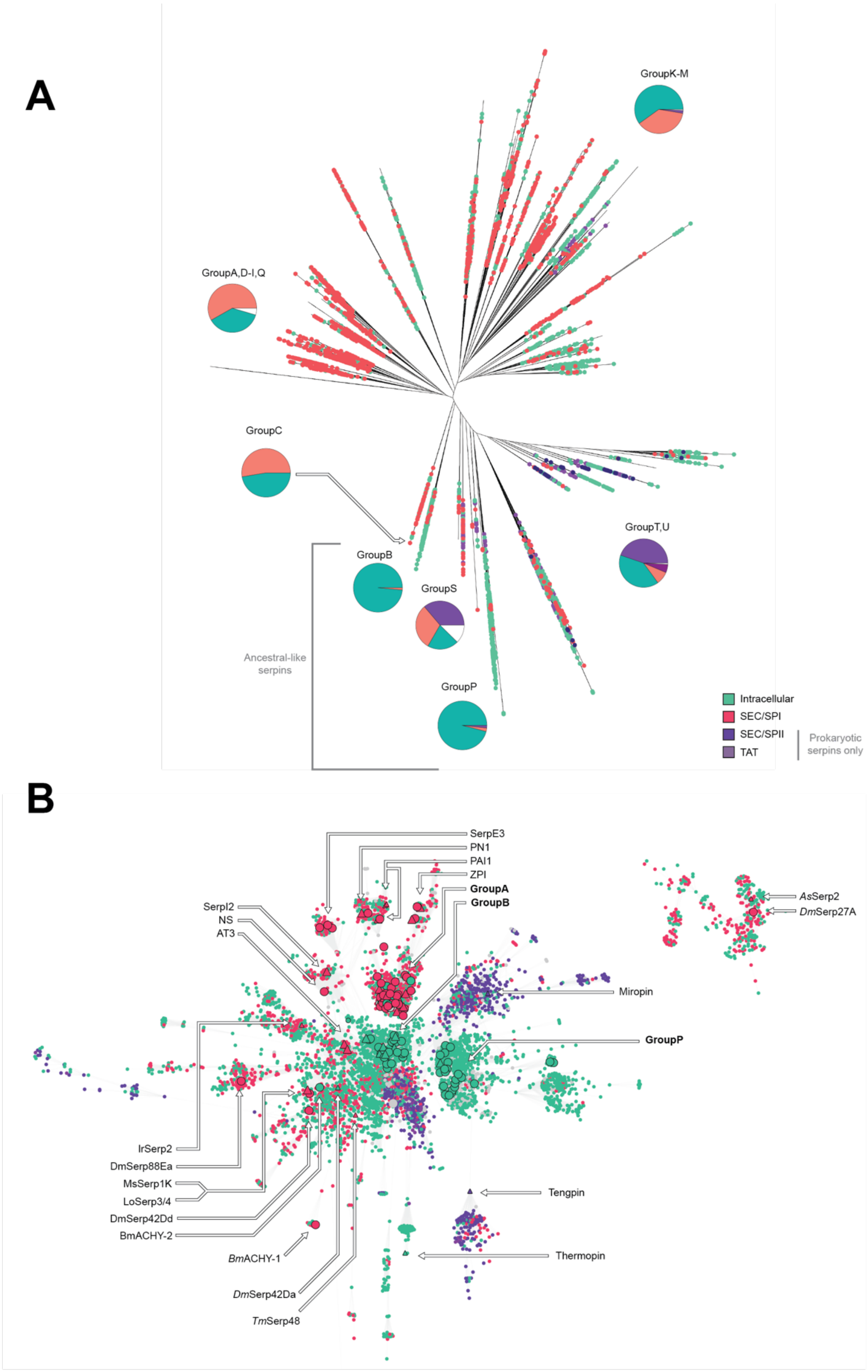
A
GroupK-M
GroupA,D-I,Q
GroupC
GroupB
GroupS
GroupT,U
GroupP
Ancestral-like serpins
Intracellular
SEC/SPI
SEC/SPII
TAT
Prokaryotic serpins only
B
SerpE3
PN1
PAI1
ZPI
GroupA
GroupB
SerpI2
NS
AT3
AsSerp2
DmSerp27A
Miropin
GroupP
IrSerp2
DmSerp88Ea
MsSerp1K
LoSerp3/4
DmSerp42Dd
BmACHY-2
BmACHY-1
DmSerp42Da
TmSerp48
Tengpin
Thermopin

**Supplementary Figure 8. Serpin superfamily ML phylogeny (A) non-redundant SSN (B) with localisation predictions**. Sequences that lack detected N-terminal transport signal peptides are highlighted in green, whereas those with transport peptides are highlighted in red and purple. Pie-charts in **A** show proportions of intracellular and extracellular serpins belonging to each major lineage in the ML phylogeny. The ancestral like eukaryotic serpins (Groups B, P) unanimously function intracellularly. The SSN hub in **B** is dominated by intracellular serpins. In both **A** and **B**, extracellular function emerged as sequences diverged from the root and hub, respectively.
